# Regulation of somatosensory mechanotransduction by Annexin A6

**DOI:** 10.1101/155812

**Authors:** Ramin Raouf, Stéphane Lolignier, Jane E. Sexton, Queensta Millet, Sonia Santana-Varela, Anna Biller, Alice M. Fuller, Vanessa Pereira, Jyoti Choudhary, Mark Collins, Stephen E. Moss, Julie Tordo, Michael Linden, John N Wood

## Abstract

Sensory neuron mechanically-activated slowly adapting currents have been linked to noxious mechanosensation. We identified a Conotoxin, Noxious Mechanosensation Blocker -1, that blocks such currents selectively and inhibits mechanical pain Using an active biotinylated form of the toxin we identified 67 binding proteins in sensory neurons and sensory neuron-derived cell lines using mass spectrometry. Annexin A6 was the most frequently identified binding protein. Annexin A6 knockout mice showed an enhanced sensitivity to mechanical stimuli. An increase in rapidly adapting currents was observed in sensory neurons alongside a decrease in slowly adapting currents. Conversely, overexpression of Annexin A6 in sensory neurons inhibited rapidly adapting currents and augmented slowly adapting currents. Furthermore, co-expression of Annexin A6 with Piezo2 led to an inhibition of Piezo-mediated rapidly adapting currents. AAV-mediated overexpression of Annexin A6 in sensory neurons attenuated mechanical pain in a mouse model of osteoarthritis. These data demonstrate a modulatory role for Annexin A6 in somatosensory mechanotransduction.

---

Pain is the greatest clinical challenge of the age in numerical terms, and mechanically-evoked pain is a major issue in many pain states, particularly those linked to arthritis and musculoskeletal pain that occur more frequently in older people ^1^. A subset of primary nociceptive neurons are directly activated by mechanical stimuli ^2^. In these neurons, mechanically-gated ion channels have been shown to be transcriptionally upregulated by NGF and their trafficking potentiated by protein kinase C ^3^. As behavioural assays of mechanical pain showed enhanced sensitivity on NGF or phorbol ester treatment, this suggests that the mechanotransducing currents described in sensory neurons in vitro have a significant role in mechanosensation ^3^. Mechanically-gated currents in sensory neurons have been classified on the basis of their adaptation kinetics ^4^. Rapidly adapting currents have been linked to proprioception and touch, via the activation of the mechanically-gated ion channel Piezo2 ^5,6^. Intermediate and slowly adapting currents are also present in sensory neuron subsets, and a link between slowly and intermediately adapting channels and noxious mechanotransduction has been made ^7^.

Although Piezo2 contributes to mechanical allodynia through EPAC mediated regulation ^8^, the principal mechanically-gated ion channels responsible for noxious mechanosensation have yet to be identified. Recently a slowly adapting mechanosensitive channel, TMEM150c or TNN, expressed in muscle spindles has been identified but its role in mechanical pain has not been explored ^9^.

A screen for toxins blocking mechanosensitive channels identified Noxious Mechanosensation Blocker-1 (NMB-1), a conotoxin peptide that selectively inhibits the persistent, non-adapting component of intermediately-adapting (IA) and slowly-adapting (SA) mechanically-activated currents in sensory neurons ^7^. NMB-1 binds to small diameter, peripherin positive neurons ^7^, known to be responsible for noxious mechanotransduction^10^. NMB-1 also inhibits mechanotransduction currents in the cochlea and blocks behavioural responses to noxious mechanical stimuli. In order to identify the targets that NMB-1 binds to, and therefore identify proteins which contribute to mechanotransduction, we developed a biotinylated form of NMB1 that retained activity. Using sensory neurons and sensory neuron derived cell lines, we performed immunoblotting and pull-down assays using biotinylated NMB-1 followed by mass spectrometry analysis. A number of candidates were identified including, as top candidate with most hits, the Ca^2+^-dependent, phospholipid-binding protein, Annexin A6. Here we describe a modulatory role for Annexin A6 in mechanotransduction in sensory neurons that can be used in vivo to block mechanical pain.

## Identification of NMB-1 binding proteins

We found that biotinylated NMB-1 (bNMB-1) can specifically bind peripherin-positive dorsal root ganglion (DRG) neurons and not large diameter sensory neurons. bNMB-1 retains its activity in blocking slowly adapting mechanically evoked currents in these neurons^7^. We used this modified form of the toxin to identify proteins that interact with NMB-1 in rat DRG neurons. We developed an immunoblotting protocol to stain interacting proteins on Western Blots of DRG or cerebellarum membrane protein extracts and found that the bNMB-1 can bind specific bands from DRG protein extracts whereas the same pattern of staining was not present in protein extracts from cerebellum (Fig. 1a). Our attempts however to identify the protein contents of the reactive bands directly from the PVDF membrane did not succeed due to low yield of protein recovery from the membrane strips (data not shown). We therefore identified binding proteins by juxtaposing alongside the blot an identically processed polyacrylamide gel stained with metallic silver to visualize all the proteins (Fig. 1a). We used LC-MS/MS spectroscopy to identify peptides present in bands from the gel that corresponded to positions of bNMB-1 reactive bands on the blot (Supplementary table 1). Using this approach, we identified a list of potential NMB-1 interacting partners. To triage this list of potential candidates and restrict it to proteins associated with the mechanotransduction complex, we used a pull-down approach and took advantage of the properties of differentiated nociceptor cell line ND-C, a DRG neuron-derived cell line that shows discrete mechanically activated currents akin to DRG neurons in culture^11,12^. We looked for bNMB-1 interacting proteins using a pull-down assay from differentiated ND-C cell protein extracts. An example of polyacrylamide gels using two different reducing conditions is shown (Fig. 1b). We sequenced the peptides present in the pull-down using LCMS/MS spectroscopy (Supplementary table 2). We reasoned that the list of proteins from the intersection of the two tables will be the smallest set containing bNMB-1 interacting proteins and found that Annexin A6 was the most abundant protein in shared between the two lists (Table 1). Annexin A6 is a calcium dependent membrane binding protein that is expressed by most sensory neurons and has been suggested to modulate Ca^2+^ conductance in DRG neurons^13^. To further confirm NMB-1 interaction with Annexin A6 in DRG neurons, we incubated WT and *Anxa6* KO DRG tissue with bNMB-1 followed by fluorescently tagged Avidin (Supplementary Fig. 1a-f). We showed a substantial (*p*=0.0298) decrease in fluorescence in *Anxa6* KO tissue (Supplementary Fig. 1g). Reduced NMB-1 binding to KO DRG neurons, together with our pull-down data, support a direct interaction of NMB-1 and Annexin A6. The residual fluorescence observed in *Anxa6* KO tissue also confirms that NMB-1 binds multiple proteins in sensory neurons.

**Figure 1.**
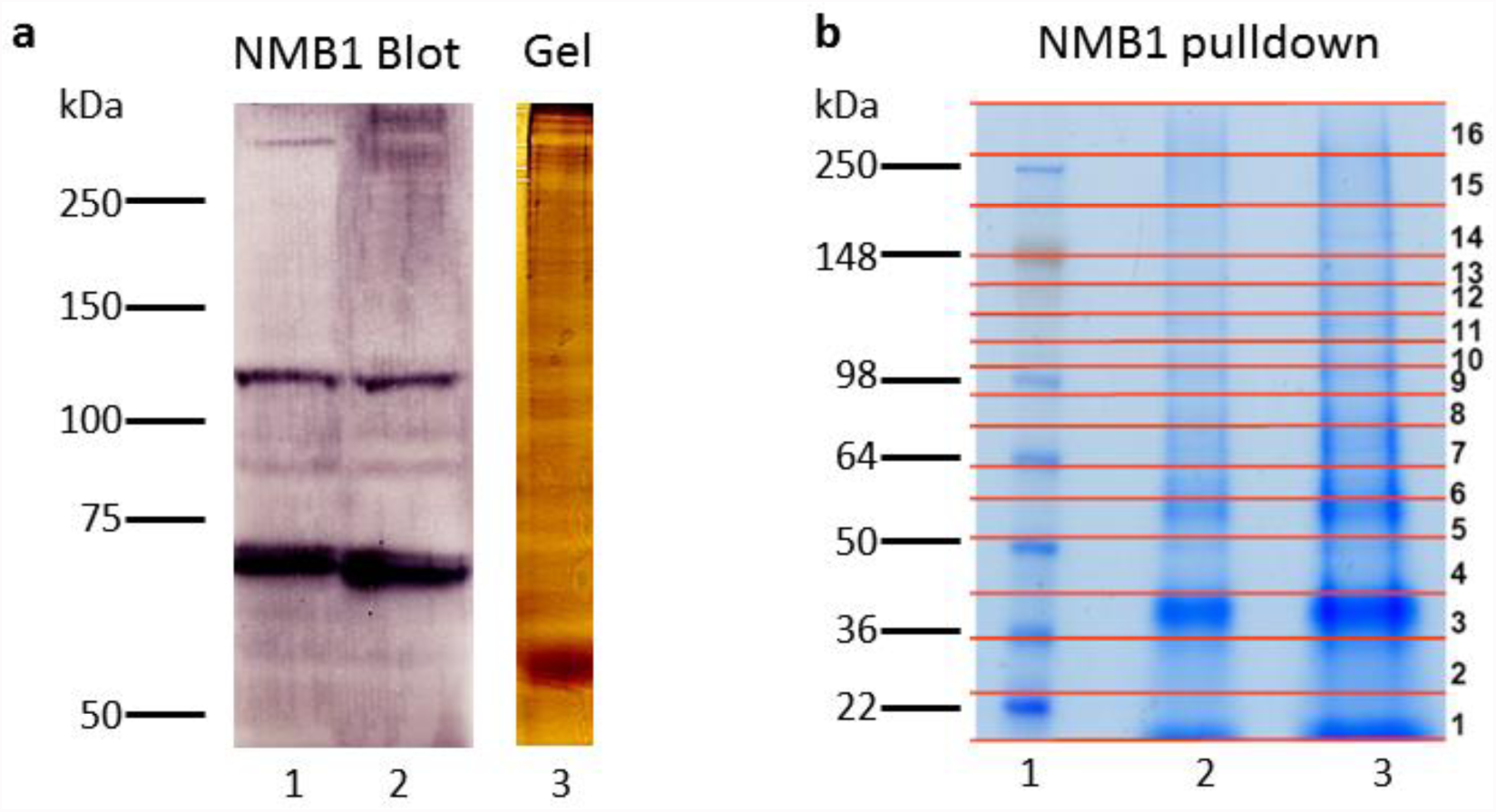
Isolation of NMB-1 peptide binding partners. a) Biotinylated NMB1 was used to visualize binding proteins on nitrocellulose blots containing cerebellum (1) or DRG (2) membrane protein extracts. A silver stained SDS gel loaded with DRG protein extract processed under the same conditions is juxtaposed alongside the NMB1 stained blot (3). Silver stained gel with reactive bands on the NMB1 blot that were excised and processed for LC MS/MS spectroscopy. b) NMB1 pulled from differentiated ND-C cell protein extracts was treated with 2% or 4% beta-mercaptoethanol and resolved on 10% SDS gel. The gel was cut into 16 strips and processed for LC MS/MS spectroscopy.

**Table 1.** NMB-1 interacting candidates shared between DRG and ND-C cells. Mouse Uniprot numbers are given. Total matches refer to combined numbers of positive peptide matches for each candidate.

| Gene name | Uniprot | Description | MW | No. matches |
| --- | --- | --- | --- | --- |
| Anxa6 | P14824 | Annexin A6, calcium-dependent phospholipid binding | 75,838 | 36 |
| Flna | Q8BTM8 | Filamin-A, actin binding | 274,142 | 28 |
| Myh9 | A2VCK1 | Myosin-9, Actin filament binding | 226,232 | 22 |
| Vim | P20152 | Vimentin, structural constituent of cytoskeleton | 53,655 | 21 |
| Anxa5 | P48036 | Annexin A5, calcium-dependent phospholipid binding | 35,730 | 8 |
| Hnrpu | O88568 | Heterogenous nuclear ribonucleoprotein U, core promoter binding activity | 87,863 | 8 |
| Snd1 | Q78PY7 | Staphylococcal nuclease domain-containing protein 1nuclease domain-containing protein 1, cadherin binding involved in cell-cell adhesion | 102,025 | 7 |
| Nono | Q99K48 | Non-POU domain-containing octamer-binding protein, chromatin binding | 54,506 | 6 |
| L1cam | Q6PGJ3 | Neural cell adhesion molecule L1, PDZ domain binding | 140,969 | 5 |
| Pdia6 | Q3THH1 | Protein disulfide-isomerase A6, protein disulfide isomerase activity, Platelet activation | 48,627 | 5 |
| Anxa4 | Q9R0V2 | Annexin A4; calcium-dependent phospholipid binding | 24,211 | 4 |
| Cpsf6 | Q6NVF9 | Cleavage and polyadenylation specific factor 6, 68kDa (Predicted), isoform CRA_b, mRNA binding | 59,116 | 4 |
| Matr3 | Q8K310 | Matrin-3, nucleotide binding | 94,572 | 3 |
| Sfxn3 | Q91V61 | Sideroflexin-3, tricarboxylate secondary active transmembrane transporter activity | 31,636 | 3 |

Based on this evidence for interaction between NMB-1 and annexin A6, we next examined the modulation of mechanotransduction in DRG neurons by annexin A6.

## Annexin A6 modulates mechanical pain and mechanotransduction in sensory neurons

Using global *Anxa6* KO mice, we investigated the role of Annexin A6 in mechanosensation *in vivo*. No difference in tactile threshold, assessed with the von Frey test, was observed between WT and *Anxa6* KO mice (50% paw withdrawal threshold of 0.64 ± 0.06 g in WT; 0.62 ± 0.10 g in *Anxa6* KO; p = 0.8386, Fig. 2a). However, in response to noxious mechanical pressure, *Anxa6* KO mice showed a 17.5% decrease in the force required to elicit a response (Fig. 2b). WT mice responded at an average of 188.8 ± 5.0 g compared to 155.8 g ± 7.7 g in KO (*p* = 0.0030). This suggests that Annexin A6 is involved in noxious, but not innocuous mechanosensation.

**Figure 2.**
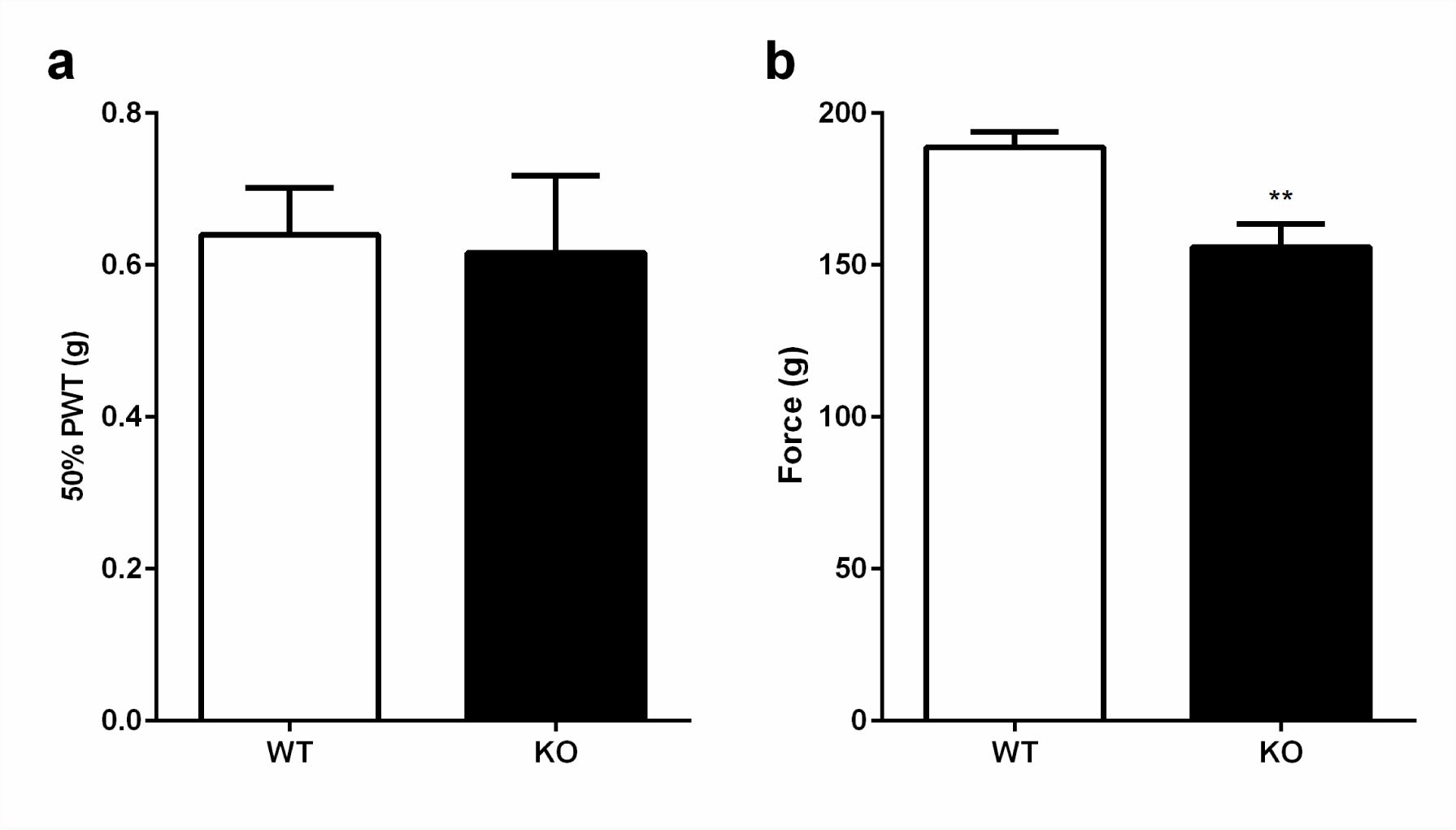
Anxa6 KO mice show increased sensitivity to noxious mechanical stimuli. (a) Responses to stimulation of the paw with von Frey hairs is unaltered in Anxa6 KO mice. (b) Anxa6 KO mice show increased sensitivity to application of a blunt probe to the tail. n = 9 ‘WT’, n = 8 ‘KO’. **p<0.01 Student’s unpaired t-test.

We then investigated mechanotransduction at the cellular level in cultured sensory neurons from WT and *Anxa6* KO mice dorsal root ganglia by recording mechanically-activated (MA) currents in whole cell patch clamp configuration ^2,14^. Currents were evoked by applying focal pressure to the cell membrane with a polished glass pipette while cells were held at -70 mV. Sensory neurons were either mechanically insensitive (non-responding, NR), or mechanically sensitive with an MA current defined by its adaptation kinetics as rapidly adapting (RA), intermediately adapting (IA), or slowly adapting (SA). Cell capacitance did not differ between WT and *Anxa6* KO neurons, whatever their response to mechanical stimulation (Fig. 3a). We observed a decrease in the percentage of neurons displaying a SA MA current, dropping from 25.6% in WT to 14.3% in *Anxa6* KO, mostly in favour of NR neurons which were increased from 20.9% in WT to 38.8% in KO (p = 0.0459; Fig. 3b). The minimum stimulation required to elicit a > 40 pA MA current was not different between WT and KO neurons, whatever the current kinetics (Fig. 3c). However, the maximum density of transient RA/IA MA currents taken together was 3.5 fold higher in *Anxa6* KO neurons compared to WT neurons (-19.7 ± 3.6 pA/pF in WT; -69.5 ± 22.1 pA/pF in KO; p = 0.047; Fig. 3d). There was a decrease in the maximum density of SA MA currents in *Anxa6* KO compared to WT neurons, although this was not significant (-85.5 ± 38.0 pA/pF in WT; -20.5 ± 8.0 pA/pF in KO; p = 0.1117). When plotted against the displacement produced by the stimulation probe, these observations show an overall increase in RA/IA MA current density and a decrease in SA MA current density in *Anxa6* KO neurons compared to WT neurons (Fig. 3e-f).

**Figure 3.**
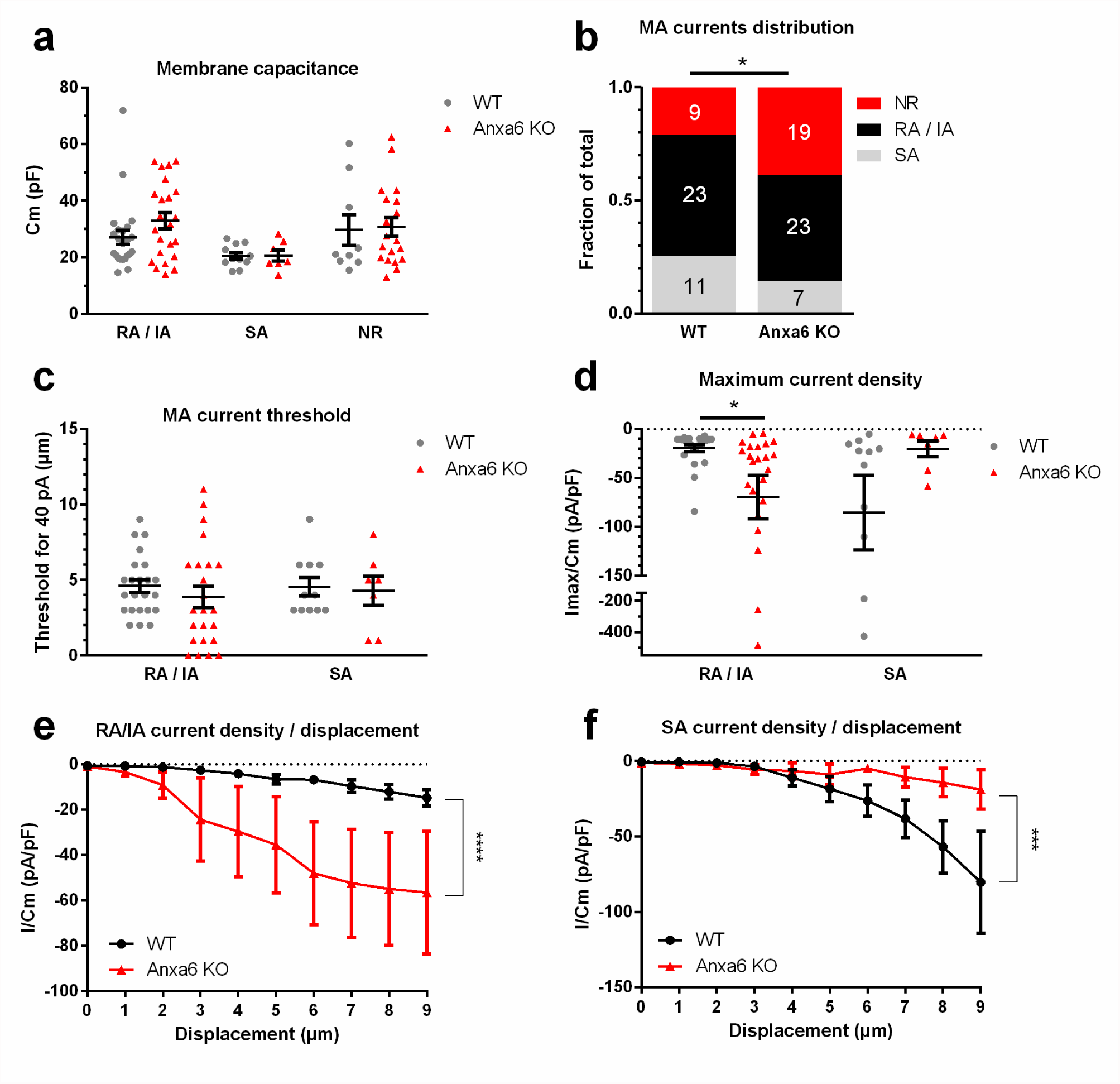
Mechanical transduction is differentially modulated in Anxa6 KO sensory neurons. Sensory neurons from WT and Anxa6 KO mice DRGs were cultured and mechanotransduction was assessed by mechano-clamp. (a) Membrane capacitances of recorded WT and Anxa6 KO neurons were not different within the 3 neuronal populations. (b) There was a larger proportion of neurons non responsive to mechanical stimulation in the Anxa6 KO group compared to WT neurons, together with a decrease in the fraction of neurons showing a SA MA current. (c) The minimal stimulation intensity required to elicit a > 40 pA MA current was not different between WT and Anxa6 KO neurons for both types of currents. (d) The maximum RA/IA MA current density recorded was higher in Anxa6 KO neurons than in WT neurons. (e) Current density plotted against the displacement of the mechanical probe shows an increase in RA/IA MA current density (two-way ANOVA, p<0.0001) and (f) a decrease in SA MA current density (two-way ANOVA, p<0.001) in Anxa6 KO neurons compared to WT neurons. (a, c, d) * p < 0.05, ** p < 0.01, *** p < 0.001 ‘WT’ vs ‘Anxa6’ by two-way ANOVA followed by Fisher’s LSD multiple comparisons test. (b) * p < 0.05 by Chi-Square contingency test. (e,f) * p < 0.05, ** p < 0.01, *** p < 0.001 two-way ANOVA; multiple comparison post tests were not significant at any given displacement. n numbers are given in (b).

We then assessed whether overexpressing Annexin A6 in DRG neurons could have the opposite effect on mechanotransduction to that induced by knocking out *Anxa6*. Cultured DRG neurons were transfected with a CMV-driven expression vector containing variant 1 of human *ANXA6* together with the fluorescent protein tdTomato, or the corresponding empty vector (EV). MA currents were recorded in tdTomato-positive neurons 2 days after transfection. Cell capacitance did not differ between cells transfected with *Anxa6* or the EV, whatever their response to mechanical stimulation (Fig. 4a). Following transfection, no change was observed in the percentage of cells expressing the different types of MA current, or NR cells (Fig. 4b). MA current thresholds were not different either between EV- and *Anxa6*-transfected cells (Fig. 4c), whatever the current kinetics, but we observed a decrease in the maximum RA/IA MA current density recorded in neurons transfected with the *Anxa6*- containing construct compared to those transfected with the EV (-96.7 ± 18.0 pA/pF within ‘EV’; -36.6 ± 11.4 pA/pF within ‘*Anxa6*’; p = 0.0060). An increase in the maximum SA MA current density recorded in neurons transfected with the *Anxa6*-containing construct was also observed, although this was not significant (-73.2 ± 23.7 pA/pF within ‘EV’; -163.8 ± 17.8 pA/pF within ‘*Anxa6*’; p = 0.0743; Fig. 4d). The significant decrease in RA/IA MA current density can be seen in current density / displacement curves (Fig. 4e) whereas for SA MA currents, neurons transfected with the EV or with the *Anxa6* expression vector showed enhanced currents (Fig. 4f).

**Figure 4.**
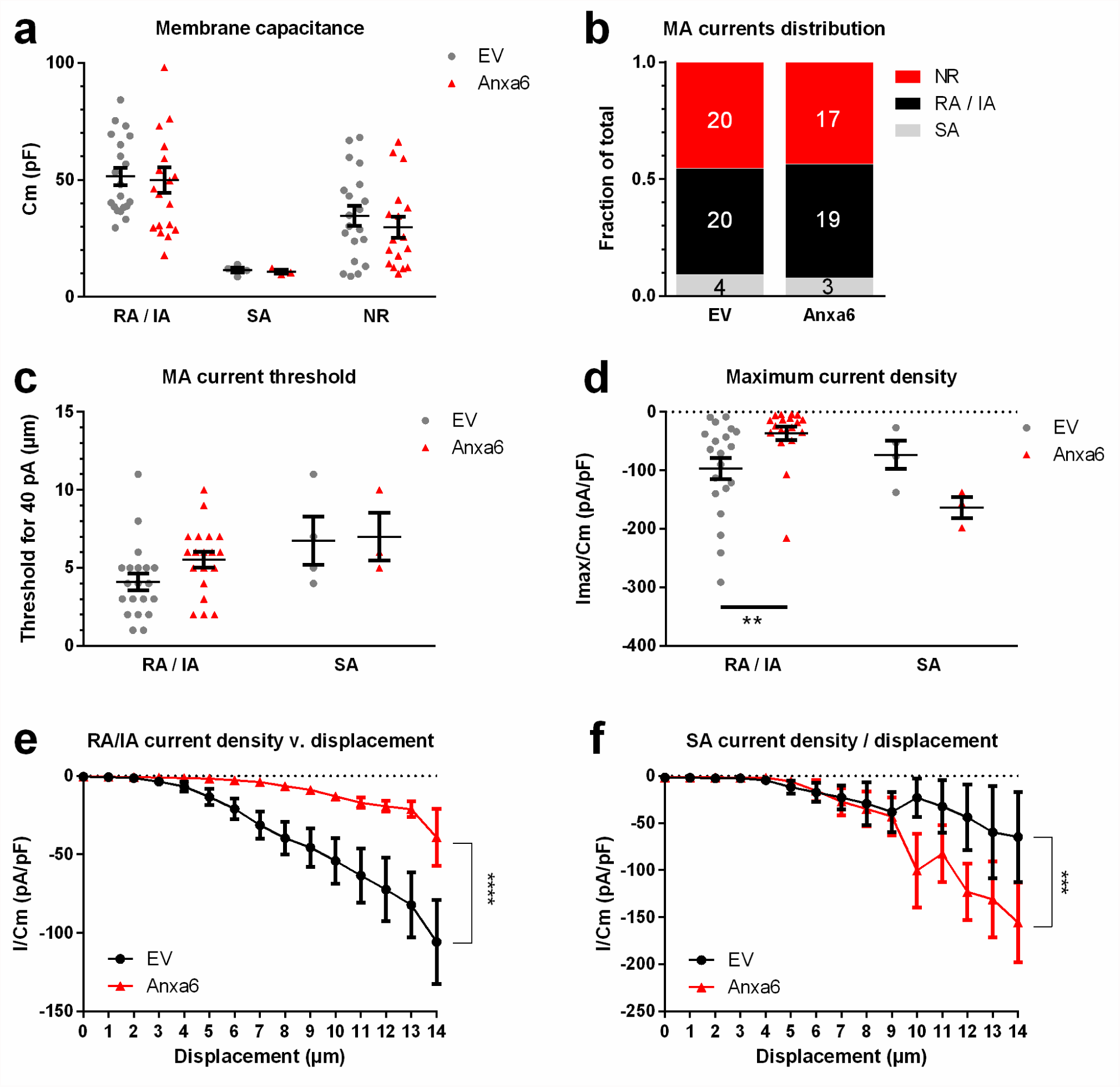
Mechanical transduction is differentially modulated in sensory neurons overexpressing Anxa6. Sensory neurons from WT mice were cultured and transfected with a human Anxa6-pIRES2-tdTomato construct or the corresponding pIRES2-tdTomato empty vector (EV). Neurons expressing tdTomato were identified through their red fluorescence and selected for mechano-clamp experiments. (a-c) Membrane capacitance (a), MA current distribution (b) and the minimal stimulation intensity required to elicit a > 40 pA MA current (c) did not differ between recorded neurons transfected either with the EV or the Anxa6-containing construct, whatever the type of MA current displayed by the cells. (d) Neurons overexpressing Anxa6 show reduced RA/IA MA current maximum density compared to control neurons. (e) Current density plotted against the displacement of the mechanical probe shows a decrease in RA/IA MA current density in neurons overexpressing Anxa6 compared to EV transfected neurons (two-way ANOVA, p<0.0001) while (f) SA MA currents are increased by Anxa6 overexpression (two-way ANOVA, p<0.001). (a, c, d) ** p < 0.01, *** p < 0.001 ‘EV’ vs ‘Anxa6’ by two-way ANOVA followed by Fisher’s LSD multiple comparisons test. (b) Chi-Square contingency test. (e,f) * p < 0.05, ** p < 0.01, *** p < 0.001 two-way ANOVA; multiple comparison post tests are not shown. n numbers are given in (b).

ANXA6 appears to be a negative regulator of transient RA/IA MA currents in DRG neurons, as shown by the downregulation of these currents in neurons overexpressing the human clone, and by their upregulation in *Anxa6* KO DRG neurons. We therefore next studied its impact on MA currents produced by PIEZO2, a mechanically gated channel responsible for low threshold, rapidly adapting, MA currents^5,15,16^. Expression vectors containing the human *Anxa6* or *Piezo2* clones (together with a tdTomato or GFP fluorescent protein, respectively), or their corresponding empty vectors, were cotransfected in ND-C cells, a DRG neuron-derived cell line that exhibits mechanosensitive currents^12^. Membrane capacitance of recorded cells was not different between the groups (Fig. 4a). *Anxa6* transfection alone did not affect the endogenous MA current threshold, maximum density or current density / probe displacement relationship (Fig. 5b-d). PIEZO2 alone produces a low-threshold current that appears to be modulated by ANXA6. Indeed, the mechanical threshold of this current increased from 2.70 ± 0.50 μm in cells transfected with *Piezo2* alone to 6.80 ± 1.32 μm in cells transfected with both *Piezo2* and *Anxa6* (p = 0.0101; Fig. 5b). The maximum density of PIEZO2 current was also reduced by 63.7% with ANXA6 co-expression (-138.0 ± 28.8 pA/pF in ‘Piezo2 + EV’; -50.0 ± 14.1 pA/pF in ‘Piezo2 + *Anxa6*’; p = 0.0160; Fig 5c). This downregulation of PIEZO2 current by ANXA6 co-expression is also significant on current density / displacement curves (Fig. 5d).

**Figure 5.**
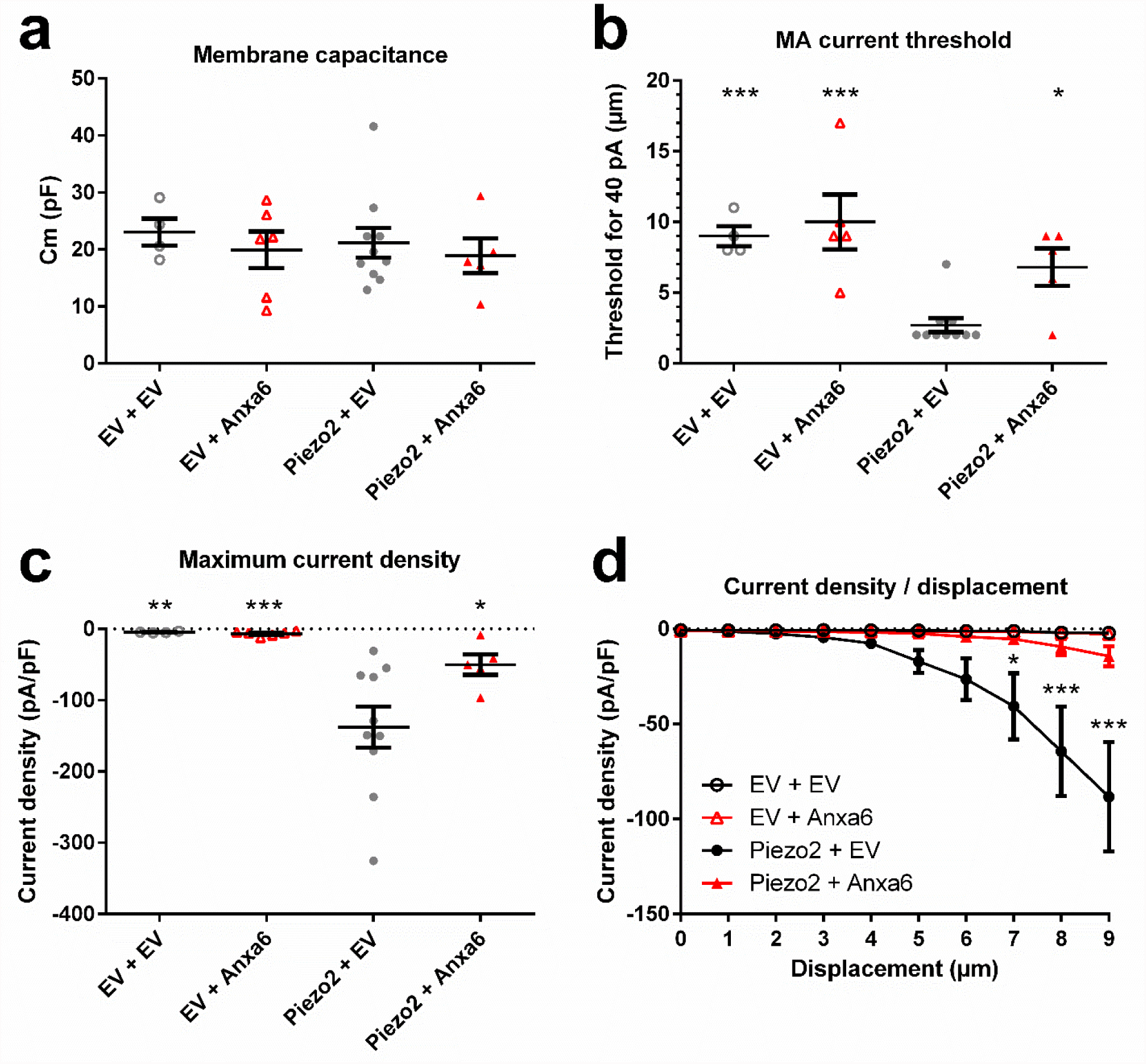
Piezo2 MA current is reduced by Anxa6 co-expression in ND-C cells. ND-C cells were cotransfected with hPiezo2-IRES-eGFP and/or hAnxa6-pIRES2-tdTomato or their respective empty vectors (EV) IRES-eGFP and pIRES2-tdTomato according to the indicated combinations, and mechanotransduction was assessed by the mechano-clamp technique. (a) The membrane capacitance of recorded cells did not differ between the groups. (b) The minimal stimulation intensity required to elicit a > 40 pA MA current was strongly reduced in Piezo2-transfected cells compared to controls. This effect is cancelled by Anxa6 co-transfection with Piezo2. (c) Co-transfection of Anxa6 with Piezo2 reduces the maximum recoded MA current density generated by Piezo2 alone by nearly 3 fold. (d) Current density plotted against the displacement of the mechanical probe shows a decrease in Piezo2 current when co-expressed with Anxa6. (a-c) * p < 0.05, ** p < 0.01, *** p < 0.001 vs ‘Piezo2 + EV’ by one-way ANOVA followed by Fisher’s LSD test. (d) * p < 0.05, *** p < 0.001 vs ‘Piezo2 + Anxa6’ by two-way ANOVA followed by Fisher’s LSD multiple comparisons test. n = 4 ‘EV + EV’, n = 6 ‘EV + Anxa6’, n = 10 ‘Piezo2 + EV’, n = 5 ‘Piezo2 + Anxa6’.

## Overexpression of *Anxa6* reduces mechanical pain in a mouse model of osteoarthritis

Having demonstrated that *Anxa6* loss of function leads to mechanical hypersensitivity *in vivo* and augmented transient RA/IA MA currents in sensory neurons *in vitro*, while *Anxa6* overexpression conversely produces a decrease in transient current density, we next assessed whether *Anxa6* overexpression in sensory neurons *in vivo* had any effects on mechanical hypersensitivity in a mouse model of osteoarthritis pain.

We generated AAV viruses (serotype TT) carrying the human *Anxa6* variant 1 clone upstream of an IRES-GFP construct and under the control of a CMV promoter. AAV-TT carrying *Anxa6,* or the empty control virus, was delivered intrathecally. Pain thresholds were followed for 12 weeks before a single intra-articular injection of monosodium iodoacetate (MIA) was given into the left knee, and behavioural assessment continued for another 3 weeks whereupon mice were sacrificed and successful transduction of DRG neurons was observed (Supplementary Fig.3). 12 weeks following AAV injection, a decrease in noxious mechanical sensitivity was seen in animals treated with the *Anxa6*-containing construct, while no difference in thermal sensitivity was observed (Fig. 6a-b). The 50% withdrawal threshold to von Frey filaments was not different between the two groups at week 12 (week 0 before MIA injection, Fig. 6c). One week after injection of MIA, all treatment groups showed a significant decrease in 50% threshold compared to the threshold recorded at baseline (Fig. 6d). Three weeks after MIA injection, control (AAV-TT-IRES-GFP treated) mice showed a further drop in 50% threshold from 0.42±0.04g (one week post MIA) to 0.27±0.03g (three weeks post MIA) while the 50% threshold of AAV-TT-*Anxa6*-IRES-GFP treated mice dropped only slightly from 0.46±0.04g (one week post MIA) to 0.41±0.06g (three weeks post MIA).

**Figure 6.**
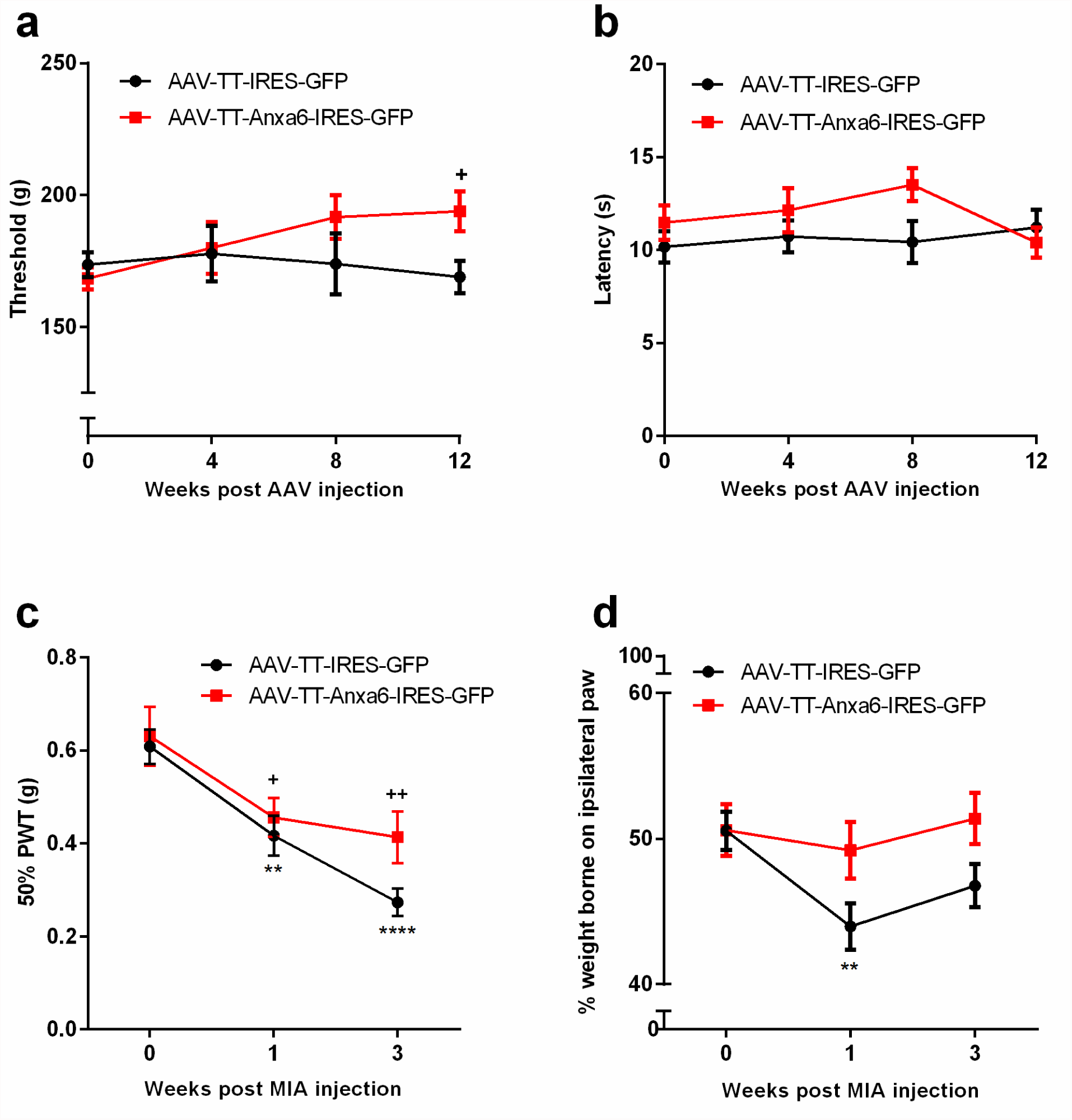
Virally mediated overexpression of Anxa6 in sensory neurons attenuates noxious mechanical sensitivity and mechanical hypersensitivity in the MIA model of osteoarthritis pain. WT mice received one intrathecal injection of AAV-TT viruses containing either a hAnxa6-IRES-GFP construct or the IRES-GFP control. Pain thresholds were then monitored for 12 weeks after which osteoarthritis was induced by a unilateral injection of monosodium iodoacetate into the knee joint and mechanical hypersensitivity was assessed for another 3 weeks. (a) 12 weeks following AAV injection, noxious mechanical sensitivity of mice injected with AAV-TT-Anxa6-IRES-GFP is significantly increased compared to pre-treatment threshold. (b) Injection of AAV-TT-Anxa6-IRES-GFP has no effect on the sensitivity of animals to noxious heat. (c) Both control and Anxa6 treated animals develop significant mechanical hypersensitivity following unilateral OA induction, though this was to a lesser extent in Anxa6 treated mice. (d) In animals treated with the control AAV-TT-IRES-GFP, MIA injection induced a significant weight bearing asymmetry, evaluated by the weight ratio between ipsilateral and contralateral rear paws. No significant change in weight bearing is observed after MIA injection in animals treated with AAV-TT-Anxa6-IRES-GFP suggesting overexpression of Anxa6 prevents onset of this weight bearing change. Data are shown as Mean±SEM; (c-f) two-way RM ANOVA with Dunnett’s multiple comparison test * p < 0.05, ** p < 0.01, *** p < 0.001 vs ‘0’ within ‘AAV-TT-IRES-GFP’; + p < 0.05, ++ p < 0.01 vs ‘0’ within ‘AAV-TT-Anxa6-IRES-GFP’; n = 12 per group. (a-b).

Attenuation of mechanical hypersensitivity in AAV-TT-*Anxa6*-IRES-GFP treated mice was also observed in the weight bearing test, designed to evaluate mechanical pain through assessing the percentage body weight applied on the MIA-injected paw. In control treated mice, weight on the ipsilateral paw dropped from 50.56±1.32% at baseline to 43.98±1.58% one week after MIA and 46.77±1.48% three weeks after MIA injection. This asymmetry was significant compared to baseline at both time points whereas *Anxa6* treated mice did not develop a significant weight bearing asymmetry at any time point. Indeed, in AAV-TT-*Anxa6*-IRES-GFP treated mice, weight on the ipsilateral paw at baseline was 50.59±1.77%; one week after MIA injection was 49.21±1.95% and three weeks after MIA injection was 51.38±1.77%. These results indicate that Annexin A6 overexpression is able to prevent the onset of the weight bearing asymmetry induced by MIA and as a proof-of-concept gene therapy approach supports a role for Annexin A6 in regulating negatively mechanosensation *in vivo*.

## Discussion

The present studies have demonstrated the ability of Annexin A6 to modulate mechanotransduction in sensory neurons. Annexins are a superfamily of Ca^2+^-dependent, phospholipid-binding proteins which are involved in diverse functional processes. They are highly evolutionarily conserved; Annexin A6, previously known as p68, calelectrin and protein III ^17^ is a compound gene which appears to derive from Annexin A5 and Annexin A10 being spliced together early in evolution. Annexin A6 is known to link the cellular cytoskeleton to the internal lipid bilayer of the cell membrane through calcium-dependent interactions with negatively charged phospholipids. We identified Annexin A6 as a binding target of the biotinylated Conotoxin NMB-1 which blocks behavioural responses to noxious mechanical stimuli and attenuates the persistent component of IA and SA MA currents. We hypothesised that Annexin A6 would therefore be a potential candidate for a role in mechanotransduction. Early studies have claimed that Annexin A6 may form a cation-selective ion channel in lipid bilayers that is a cGMP sensor^18^. Other studies have suggested that low pH potentiates the ability of Annexin A6 to form cation channels^19^.

We found that Annexin A6 KO mice have heightened sensitivity to noxious mechanical stimuli and sensory neurons from these mice show increased transient, RA and IA, mechanically evoked currents. Conversely, when overexpressing Annexin A6 in a mechanosensitive cell line, we were able to dramatically reduce the current produced by Piezo2, the well-defined mechanotransduction channel. Similarly, overexpression of Annexin A6 in sensory neurons reduced the amplitude of RA and IA currents which, together, demonstrate that Annexin A6 is able to regulate MA currents *in vitro*. Thus Annexin A6 has a general role in negatively regulating mechanotransduction currents. This data provides the first evidence of a potential role for Annexin A6 in cutaneous mechanosensation. Interestingly, Annexin A6 has been linked to roles in mechanosensory functions in other physiological systems. For example, in mice overexpressing Annexin A6 in the heart, there is a decrease in the contractility of cardiomyocytes, presumed to be the result of reduced free intracellular Ca^2+^ and reduced Ca^2+^ release upon depolarisation ^20^. Similarly, in skeletal muscle, Annexin A6 regulates the gating properties of the sarcoplasmic reticulum Ca^2+^ release channel, potentially linking it to a role in excitation-contraction coupling responsible for muscle contraction ^21^.

Current understanding of the functions of Annexin A6 provides insight into the potential mechanisms by which it could regulate mechanotransduction. In conditions of low Ca^2+^ annexins are soluble monomeric proteins though when intracellular Ca^2+^ levels rise, annexins can bind to phospholipid membranes ^13^. It is proposed that to bind the membrane, annexins form trimers and aggregate into a hexagonal array surrounding a target protein ^13,22,23^. This is known to affect membrane properties, for example altering Ca^2+^ mediated membrane fluidity ^24-27^ and potentially interfering with protein-protein or protein-phospholipid interactions as well as affecting Ca^2+^ dependent second messenger pathways ^28^. If Annexin A6 were able to regulate membrane fluidity, increasing membrane rigidity in the same way as Annexin A5 for example ^27^, it is possible that this could increase the force required to elicit a mechanically activated response. In this situation, it is possible that RA and IA currents, which are typically evoked by a lower intensity of mechanical displacement, would be most sensitive to changes produced by overexpressing Annexin A6 in cells. This is consistent with our overexpression data in DRG where we see an attenuation of RA and IA current amplitude. Whether Annexin A6 expression is regulated in sensory neurons in physiological conditions is currently unknown but evidence from cardiomyocytes demonstrates the potential for its expression to be dynamically regulated in the cell^29^.

Our gene therapy data suggests that Annexin A6 mediated regulation of mechanosensation also has the potential to be clinically useful. Gene therapy offers a target specific alternative to existing pharmacological interventions and has already been shown to be a viable treatment method for analgesia in the clinic^30^. Similarly, in preclinical models, gene therapy for pain relief is effective in multiple pain conditions^31-35^ including rheumatoid arthritis^36^. We used a novel AAV serotype which has a high tropism for neurons, AAV-TT, to overexpress human ANXA6 *in vivo* and found that overexpressing ANXA6 is sufficient to attenuate mechanical hypersensitivity in the MIA model of osteoarthritis pain.

In addition to Annexin A6, we tested another peptide binding partner of NMB-1 for a role in mechanosensation. *ATP6V182* is the brain isoform of a vacuolar ATP synthase subunit which is expressed in cochlear hair cells and has been linked to a role in hearing^37^. However, expression of *ATP6V182* in HEK293T cells was insufficient to produce a mechanically activated current (Supplementary Fig.2 a-b) though co-expression of the subunit with Piezo2 partially reduced its peak amplitude in a similar way to Annexin A6 (Supplementary Fig. 2a-b). Other studies have found that some proteins with apparent roles in mechanosensation, such as TRPC3, TRPC6 and TRPA1, appear to function in a context dependent manner^38,39^. Nonetheless, expression of *ATP6V182* in a neuronal cell line with endogenous mechanosensitivity did not produce an MA current (Supplementary Fig. 2c-d) suggesting that the subunit has either an indirect or no role in mechanotransduction. Other candidate proteins identified by NMB-1 peptide binding are being examined as potential mechanotransducers. It is becoming increasingly apparent that, as with redundant mechanisms of thermosensation, mechanotransduction and mechanosensory function as a whole is likely to involve multiple proteins and mechanisms.

## Methods

### Ligand blotting assay using DRG and cerebellum membrane protein extracts

Membrane proteins were isolated using a previous published protocol^40^. Briefly, DRG (70 ganglia) and cerebellum were excised from p0 rats and homogenized using a glass Teflon homogenizer in homogenization buffer (0.32 m sucrose, 10 mm HEPES, pH 7.4, and 2 mm EDTA plus protease inhibitors leupeptin, pepstatin, phenylmethylsulfonyl fluoride, benzamidine, aprotinin). The homogenates was centrifuged for 15 min at 1000 × *g* and the supernatant centrifuged at 200,000 × *g* for 15 min and extracted with lysis buffer 50 mm HEPES, pH 7.4, 2 mm EDTA, protease inhibitors, 4% SDS and 2% Triton X-100. The samples were resolved using SDS-PAGE on 8 - 12% gels and transferred to PVDF or nitrocellulose filters (R&D systems). The membrane was blocked with 0.1% Tween-20 in PBS, pH7.4 (wash buffer) and incubated with biotinylated NMB-1 (2 or 5 μM) for 15 min. The membranes were washed and treated with chilled 4% paraformaldehyde. This crosslinking step was found to enhance binding of NMB-1^7^. The membranes were incubated with alkaline phosphatase-conjugated streptavidin (Chemicon) for 2 hours at room temperature. bNMB-1 was detected using alkaline phosphatase colour reaction using BCIP/NBT kit from Vector Laboratories according to the manufacturer’s instructions. Levamisole Solution (Vector Laboratories) was added to inhibit endogenous alkaline phosphatase activity. The appearance of blue precipitate was monitored on both bNMB-1 treated and control blots. Control experiments were also carried out where NMB-1 was substituted with goat anti biotin IgG to ensure specificity of NMB-1 blotting assay. Reactive bands on the PVDF or nitrocellulose filters were cut and sent to MRC Clinical Sciences Centre (Imperial College, London UK) for analysis. However, attempts to extract the proteins from the filter and process for LC-MS/MS or perform in situ digestion did not produce sufficient quantities of material required for mass spectroscopy. As an alternative approach, we split into two a mini-gel after resolving proteins samples. One half was electrotransferred onto nitrocellulose filter and processed for bNMB-1 binding; and the other half was stained with metallic silver using Silver Stain Kit (Pierce, ThermoFisher). The silver stained half was aligned with the bNMB-1 blot based on protein markers’ position and other landmarks. Gel bands corresponding to the position of the reactive bands on the blots were excised from the gel. The samples were analyzed using LC-MS/MS on an Agilent Q-TOF LC/MS.

### NMB-1 pull-down from ND-C cell extracts

Seven 10 cm plates of sub-confluent, actively dividing ND-C cells were treated for three days with D-MEM containing 1 mM dibromo-cAMP and 0.2% FBS. The cultures were washed with PBS and incubated with 5 μM biotinylated NMB-1 for 3.5 minutes. After wash, chilled 4% PFA was added to cells for 10 minutes and washed. Extraction buffer containing protease inhibitors, 0.1% SDS and 1% Triton X-100 was added directly to the plate. The cells were scraped and extracted for three hours on ice. The protein extract was vortexed, spun down and incubated with prepared Neuroavidin gel matrix (ThermoFisher Scientific) overnight at 4 C. The beads were washed three times and the bound proteins eluted according to the manufacturer’s instructions by heating to 95 C for 10 minutes in 2% or 5% β–mercaptoethanol. Control samples were processed as above omitting biotinylated NMB-1.

Eluates from NMB-1 columns were separated by SDS-PAGE electrophoresis using a NuPAGE 4-12% Bis-Tris gel (1.5 mm × 10 well, Invitrogen) using MOPS buffer. The gel was stained overnight with colloidal Coomassie blue (Sigma). Each lane was excised into 16 bands that were cut into 1-2 mm cubes and in-gel digested overnight using trypsin (sequencing grade; Roche). Peptides were extracted from gel bands twice with 50% acetonitrile/0.5% formic acid and dried in a SpeedVac (Thermo). Peptides were resuspended using 0.5% formic acid were analysed online using an Ultimate 3000 Nano/Capillary LC System (Dionex) coupled to an LTQ FT hybrid mass spectrometer (Thermo Electron) equipped with a nanospray ion source. Peptides were desalted on-line using a micro-Precolumn cartridge (C18 Pepmap 100, LC Packings) and then separated using a 30 min RP gradient (4-32% acetonitrile/0.1% formic acid) on a BEH C18 analytical column (235 μm id × 10 cm) (Waters).

The mass spectrometer was operated in standard data dependent acquisition mode controlled by Xcalibur 2.0. The instrument was operated with a cycle of one MS (in the FTICR cell) acquired at a resolution of 100,000 at m/z 400, with the top five most abundant multiply-charged ions in a given chromatographic window subjected to MS/MS fragmentation in the linear ion trap.

### LC-MS/MS analysis of NMB-1 binding proteins

Elutions from NMB-1 pull-down experiments were separated by SDS-PAGE electrophoresis using a NuPAGE 4-12% Bis-Tris gel (1.5 mm × 10 well, Invitrogen) using MOPS buffer. The gel was stained overnight with colloidal Coomassie blue (Sigma). Each lane was excised into 16 bands that were cut into 1-2 mm cubes and in-gel digested overnight using trypsin (sequencing grade; Roche). Peptides were extracted from gel bands twice with 50% acetonitrile/0.5% formic acid and dried in a SpeedVac (Thermo). Peptides were resuspended using 0.5% formic acid were analysed online using an Ultimate 3000 Nano/Capillary LC System (Dionex) coupled to an LTQ FT hybrid mass spectrometer (Thermo Electron) equipped with a nanospray ion source. Peptides were desalted on-line using a micro-Precolumn cartridge (C18 Pepmap 100, LC Packings) and then separated using a 30 min RP gradient (4-32% acetonitrile/0.1% formic acid) on a BEH C18 analytical column (235 μm id × 10 cm) (Waters).

The mass spectrometer was operated in standard data dependent acquisition mode controlled by Xcalibur 2.0. The instrument was operated with a cycle of one MS (in the FTICR cell) acquired at a resolution of 100,000 at m/z 400, with the top five most abundant multiply-charged ions in a given chromatographic window subjected to MS/MS fragmentation in the linear ion trap. All data were processed using BioWorks V3.3 (Thermo Electron) and searched using Mascot server 2.2 (Matrix Science) against a mouse IPI sequence database (June, 2007) using following search parameters: trypsin with a maximum of 2 mis-cleavages, 20 ppm for MS mass tolerance, 0.5 Da for MS/MS mass tolerance, with 3 variable modifications of Acetyl (Protein N-term), Carbamidomethyl (C), and Oxidation (M). False discovery rates determined by reverse database searches and empirical analyses of the distributions of mass deviation and Mascot Ion Scores were used to establish score and mass accuracy filters. Application of these filters to this dataset resulted in a < 1% false discovery rate as assessed by reverse database searching. Protein hits from all datasets were BLAST-clustered using a threshold of 95% sequence homology over at least 50% of sequence length.

## Proteomics data processing

All data were processed using BioWorks V3.3 (Thermo Electron) and searched using Mascot server 2.2 (Matrix Science) against a mouse IPI sequence database (June, 2007) using following search parameters: trypsin with a maximum of 2 mis-cleavages, 20 ppm for MS mass tolerance, 0.5 Da for MS/MS mass tolerance, with 3 variable modifications of Acetyl (Protein N-term), Carbamidomethyl (C), and Oxidation (M). False discovery rates determined by reverse database searches and empirical analyses of the distributions of mass deviation and Mascot Ion Scores were used to establish score and mass accuracy filters. Application of these filters to this dataset resulted in a < 1% false discovery rate as assessed by reverse database searching. Protein hits from all datasets were BLAST-clustered using a threshold of 95% sequence homology over at least 50% of sequence length.

### Animals

All experiments were approved by the UK Home Office and performed under a Home Office project licence (PPL 70/7382). 7-8 weeks old male C57Bl6/J mice, *Anxa6* knock-outs or wild-type littermates were used and kept under 12/12 light cycle with food and water *ad libitum*. *Anxa6* global knock-outs were previously generated by disruption of exon 3 with insertion of a neo cassette^41^. Experimenters were blind to genotype or treatment of the animals. AAV viruses were injected intrathecally (10 μl at 2.4 × 10^12^ VP/ml) 20 min following intravenous injection of mannitol (200 μl at 25%) to potentiate sensory neurons transduction^42^. Osteoarthritis was induced by injection of 0.5 mg monosodium iodoacetate (Sigma, UK) into the right knee joint cavity under light isoflurane anaesthesia^43^.

### Plasmids

hANXA6 transcript variant 1 clone (NP_001146) was purchased from Origene (cat. #RC202086) and subcloned into a pIRES2-tdTomato vector, originally made from a pIRES2-AcGFP1 backbone. hPiezo2 (NP_071351 with SNPs rs7234309 and rs3748428) was cloned into an IRES-eGFP vector as previously described^8^.

### AAV-TT virus production and purification

HEK293T cells were co-transfected with a pAAV-CMV-h*ANXA6*-IRES-GFP plasmid containing the hAnxa6 transcript variant 1 clone (NP_001146) upstream a GFP, flanked by the ITRs of AAV2, together with a helper plasmid providing the *rep* and AAV-TT *cap* genes in trans^44^. 72 h after co-transfection with PEI-max (Polysciences, 3.5 ml per mg DNA), cells were harvested by centrifugation and lysed by 4 freeze-thaw cycles (from -80 to 37°C) in lysis buffer (150 mM NaCl, 50 mM Tris pH 8.5) before benzonase treatment (5000 U / cell factory 10 chamber, corresponding to 6320 cm2 growth area, 37°C for 30 min). Lysate was cleared by centrifugation and filtered at 0.22 μm. Cell culture supernatant was also harvested and precipitated by addition of ammonium sulfate to 31.3%. The resulting pellet was also resuspended in lysis buffer, treated with benzonase, clarified by centrifugation and filtered.

Recombinant AAV-TT virions were purified by fast protein liquid chromatography (ÄKTApurifier system, GE Healthcare) and an AVB sepharose affinity column (buffer A: PBS, pH 8; buffer B: 0.5 M glycine, pH 2.7). The collected fractions were then dialysed against PBS overnight and viruses were concentrated with Vivaspin 20 centrifugal concentrators (100 kDa MWCO, Sartorius). Viral genomes titer as well as capside titer was determined by real time quantitative PCR and SDS-PAGE, respectively. Primers used for PCR quantification were the following: GFP forward 5’ GAC GGC AAC ATC CTG GGGCAC AAG 3’ and GFP reverse 5’ CGG CGG CGG TCA CGA ACT C 3’. A standard curve was realized by loading known quantities of the linearized pAAV-CMV-AnxA6-IRES-GFP vector plasmid. The vectors were finally diluted to 2.4 × 10^12^ VP/ml in saline before injection.

### Behaviour

Tactile sensitivity of the plantar surface of animals was measured using the up-down method for obtaining 50% threshold using von Frey hairs^45^. Animals were habituated to the von Frey chambers for 2 h before the experiment. Noxious mechanical sensitivity was assessed using Randall Selitto apparatus. Briefly, a blunt probe was applied to the tail with increasing force and the force at which the animal withdrew was recorded^46^. The test was performed 3 times for each animal.

Animals’ weight distribution was assessed using an incapacitance meter (Linton Instrumentation). Mice were placed in rearing position, with the upper body in a cylindrical chamber. The average weight applied by each hindpaw on the two sensors was measured over a 5s period of steady behaviour with both paws in contact with the sensors. Ipsilateral/contralateral weight ratios presented were calculated from 3 averaged consecutive measurements.

### Immunohistochemistry

DRG and spinal cord were taken from mice following intracardiac perfusion with 4% paraformaldehyde (PFA), post-fixed for 2 h in 4% PFA and cryoprotected overnight in 30% sucrose (all solutions made from phosphate buffer saline). Tissues were embedded in OCT compound and stored at -20°C until 10 and 30 μm cryosections of DRGs and spinal cord, respectively, were performed and mounted onto Superfrost Plus slides (Thermo Scientific). Slides were dried for 2 h, blocked in 10% goat serum, and incubated overnight with a primary antibody anti-GFP (Abcam #ab290, 1:2000), anti-Anxa6 (Abcam #ab31026, 1/400), and/or with biotin-conjugated IB4 (Sigma #L2140, 1:500). Slides were then incubated 2 h with Alexa Fluor 488-conjugated goat anti-rabbit IgG (Abcam #ab150077, 1:1000) and Alexa Fluor 568-streptavidin conjugate (Life technologies, #S-11226, 1:1000). For NMB1/Annexin A6 interaction studies, lumbar DRG tissue was extracted from n=3 mice per genotype and snap frozen on dry ice in OCT before being stored at -80°C. Sections were prepared as described above before being incubated in ice cold 4% PFA for 8 mins. Samples were blocked with protein block (Abcam #64226) for 2 hours at RT before being incubated with 2μM bNMB-1 (or no NMB-1 as control) in protein block overnight at 4°C. Samples were then treated with FITC-conjugated Avidin (Invitrogen #434411; 1:1000) at RT for 2 hours. Images were taken on a Leica SP8 confocal microscope. Fluorescence was quantified in 6-10 sections per mouse using ImageJ software and normalized to background fluorescence.

## Cell culture and electrophysiology

### Culture and transfection of DRG neurons

Immediately after CO_2_ euthanasia, thoracic and lumbar dorsal root ganglions were removed from 6-12 weeks old mice and subsequently digested for 35 min at 37°C in Ca^2+^- and Mg^2+^-free Hanks’ balanced salt solution (HBSS) containing 5 mM 4-(2-hydroxyethyl)-1-piperazineethanesulfonic acid (HEPES), 10 mM glucose, 5 mg/ml collagenase type XI and 10 mg/ml dispase. After digestion, ganglions were gently triturated in Dulbecco’s modified Eagle’s medium (DMEM) containing 10% qualified foetal bovine serum (FBS) using fire-polished glass Pasteur pipettes. When required, neurons were electroporated using the Neon Transfection System (Life Technologies) according to the supplier’s recommendations, in 10 μl tips and with 0.6 μg plasmid per reaction, applying 2 × 20 ms pulses of 1100 V. Cells were plated on poly-L-lysine and laminin coated dishes, in DMEM containing 10% FBS and 125 ng/ml 7S nerve growth factor (NGF). Neurons were kept at 37°C in 5% CO_2_ and used for patch clamp studies the following two days, or at 48 ± 4 h for electroporated neurons.

### Culture and transfection of ND-C cells

ND-C cells, a hybridoma between neonatal rat dorsal root ganglia neurons and mouse neuroblastoma^11^, were cultured in DMEM containing 10% FBS and kept at 37°C in 5% CO_2_. Two days prior to patch clamp recordings, the cells were electroporated using the Neon Transfection System according to the supplier’s recommendations, in 10 μl tips and with 0.6 μg plasmid per reaction, applying 3 × 20 ms pulses of 1100 V.

### Electrophysiological recordings

Small (< 30 pF) or large (> 30 pF) DRG neurons whose somata were not in contact with those of other neurons, and who were displaying clear fluorescent signal when transfection was performed, were selected for electrophysiological recordings. Patch pipettes were pulled from borosilicate glass capillaries using a PC-10 puller (Narishige Group) and had a resistance of 1.5-3.5 MΩ. The pipette solution contained (in mM): 125 CsCl, 4.8 CaCl_2_, 1 MgCl_2_, 4 MgATP, 0.4 Na_2_GTP, 10 ethylene glycol tetraacetic acid (EGTA) and 10 HEPES (pH: 7.4 adjusted with CsOH; osmolarity: 310 mOsm adjusted with sucrose). The bath solution contained (in mM): 132 NaCl, 3 KCl, 2.5 CaCl_2_, 1 MgCl_2_, 10 glucose, 10 HEPES (pH: 7.4 adjusted with NaOH; osmolarity: 310 mOsm adjusted with sucrose).

Voltage and current clamp recordings were performed using a MultiClamp 200B amplifier and an Axon DigiData 1440A digitizer (Molecular Devices). Data were recorded and stored using Clampex 10 (Molecular Devices). Recordings were low-pass filtered at 5 kHz and sampled at 20 kHz. Capacity transients were cancelled; however, series resistances were not compensated. Voltages were not corrected for junction potentials and recordings were performed at room temperature. Holding command was set at -70 mV before and during mechanical stimulation. When required, Vm measurements were performed immediately after achieving whole cell configuration. Off-line analysis and fits were performed using Clampfit 10 (Axon Instruments, Molecular Devices Inc.).

## Mechanosensory current analysis

Mechanical stimulation of cell bodies was achieved using a heat-polished glass pipette (tip diameter of approximately 2 μm) controlled by a piezo-electric crystal drive (Burleigh LSS-3000) and placed above the centre of the cell soma at an angle of 70^°^ to the surface of the dish. The probe was positioned so that a 10 μm movement did not visibly contact the cell but that an 11 μm movement, considered as a 1 μm stimulus, produced an observable membrane deflection. The probe was moved at a speed of 1 μm/ms. 250 ms mechanical steps were applied every 10 s in 1 μm increments.

Currents were characterized as a function of the signal given by the highest stimulation intensity applied before the seal was lost. Cells were considered as non-responding when no current > 20 pA was observed in response to a 12 μm stimulus. Responding cells were then classified regarding to the adaptation kinetics of their currents. Rapidly adapting MA currents had a decay kinetic that was best described by a bi-exponential fit. Intermediately adapting MA currents had a decay kinetic that was best described by a mono-exponential fit. Slowly adapting currents showed minimal decay (<20%) during the 250ms stimulation.

## Data analysis

Raw data were analysed for statistical significance using either unpaired Student’s t-test, one-way or two-way ANOVA with repeated measures, as appropriate, followed by Dunnett’s multiple comparison test, Fisher’s LSD multiple comparisons test, or Chi-Square contingency test as indicated. Analysis were performed using GraphPad Prism 6 software. All values are shown as mean ± SEM.

## Contributions

R.R, S.L and J.N.W. designed the study. R.R performed NMB-1 experiments and mass spectrometry analysis. J.C and M.C performed mass spectrometry. S.L. carried out electrophysiological studies and gene therapy studies. J.E.S performed behavioural assays and gene therapy studies. Q.M bred mice and Q.M and S.S.V performed behavioural studies. A.B, and V.P contributed to molecular and gene therapy experiments. A.M.F performed ATP6V1B2 electrophysiological studies. S.E.M provided transgenic mice. J.T and M.L provided and grew virus. R.R, S.L, J.E.S and J.N.W. wrote the paper, to which all authors contributed.

## Acknowledgements

We thank Arthritis Research UK and the Wellcome Trust for their generous support.

